# Visual field asymmetries in responses to ON and OFF pathway biasing stimuli

**DOI:** 10.1101/2024.07.15.603635

**Authors:** Martin T.W. Scott, Alexandra Yakovleva, Anthony Matthew Norcia

## Abstract

Recent reports suggest the ON and OFF pathways are differentially susceptible to selective vision loss in glaucoma. Thus, perimetric assessment of ON- and OFF-pathway function may serve as a useful diagnostic. However, this necessitates a developed understanding of normal ON/OFF pathway function around the visual field and as a function of input intensity. Here, using electroencephalography, we measured ON- and OFF-pathway biased contrast response functions in the upper and lower visual fields. Using the steady-state visually evoked potential paradigm, we flickered achromatic luminance probes according to a saw-tooth waveform, the fast-phase of which biased responses towards the ON or OFF pathways. Neural responses from the upper and lower visual fields were simultaneously measured using frequency tagging - probes in the upper visual field modulated at 3.75Hz, while those in the lower visual field modulated at 3Hz. We find that responses to OFF/decrements are larger than ON/increments, especially in the lower visual field. In the lower visual field, both ON and OFF responses were well described by a sigmoidal non-linearity. In the upper visual field, the ON pathway function was very similar to that of the lower, but the OFF pathway function showed reduced saturation and more cross-subject variability. Overall, this demonstrates that the relationship between the ON and OFF pathways depends on the visual field location and contrast level, potentially reflective of natural scene statistics.

## Introduction

Differences in psychophysical performance, physiology, and anatomy have been reported between the upper and lower visual fields (the UVF and LVF, respectively). Examples of a performance advantage in the LVF come from studies of contrast sensitivity (Abrams et al., 2012; Cameron et al., 2002), hue discrimination (Levine & McAnany, 2005), motion and shape perception (Zito et al., 2016) (to name a few, see Himmelberg et al., 2023 for a review). As early as in the retina, an LVF bias has been observed in the density of midget retinal ganglion cells (Curcio & Allen, 1990), and in human cortex, the LVF is represented by a disproportionately large amount of V1 (Benson et al., 2021). A potential contribution from the cortex is supported by recent detailed work which simulated known VF asymmetries in retinal structure and function in a computational observer (Kupers et al., 2022). Despite the addition of a simplified retinal ganglion cell (RGC) model, Kupers and colleagues’ simulation did not produce the behavioral VF asymmetries observed in humans. This result implies that retinal asymmetry is enhanced by cortex, although not all RGCs in the retina were included in their computational observer model.

A prominent bifurcation of the RGCs is their division into ON and OFF subdivisions, which encode local increments and decrements in light (respectively) (Kremers et al., 1993). The ON- and OFF-pathways remain segregated at the primate lateral geniculate nucleus (Reid & Shapley, 1992), and are the building blocks of cortical simple-cell ON and OFF sub-regions. In the absence of functional asymmetry, a parallel encoding of local increments and decrements alone is advantageous as it reduces metabolic cost while preserving informational capacity (Gjorgjieva et al., 2014). Indeed, early work considered the ON and OFF subdivisions to be functionally symmetric (Schiller, 1992). However, there is a growing corpus indicating that the human ON and OFF pathways are not symmetric; both from psychophysics (Bowen et al., 1989; Komban et al., 2011) and electrophysiology (Norcia et al., 2020; Zemon & Gordon, 2006; Zemon et al., 1988). It should be noted that ON-pathway & OFF-pathway symmetry is implicitly assumed when experimenters use an unsigned definition of luminance contrast, as many do. Unsigned contrast definitions (like Michelson contrast) are often used for stimuli with periodic spatial patterns, like gratings or checker-boards. They do not distinguish between a dark element on a grey background, and a light element on a grey background (while ON- and OFF-RGCs do). The Weber contrast is a signed definition of contrast that retains this contrast polarity, making it more useful for probing the ON and OFF pathways. See Westheimer (2007) for a historical perspective on the dominance of unsigned contrast. Crucially, if the ON and OFF pathways are differently tuned, it is possible that this tuning interacts with visual field location and contributes to VF performance asymmetries. Describing the degree of ON- and OFF-pathway VF asymmetry may then allow us to better simulate human behavior. Additionally, it would provide a more detailed understanding of the perceptual experiences of individuals with pathology that affects one pathway more than the other, like amblyopia (Pons et al., 2019) and glaucoma (Norcia et al., 2022).

Why might we expect ON- and OFF-pathway spatiotemporal tuning to vary by visual field location? One driving force for tuning asymmetry is an asymmetric distribution of information in natural scenes. Neurons have a limited range of response magnitudes, and large magnitude responses are metabolically costly. So, it is metabolically efficient to represent the most commonly encountered contrasts at low response magnitudes and with greater granularity (i.e., using a sigmoidal non-linearity). The notion of allocating neural sensitivity based on the frequency of occurrence is a form of “tuning” central to information theory (Barlow et al., 1987). If information theory holds, biases in contrast sensitivity & discriminability across the visual field can be predicted by spatial biases in contrast information. Interestingly, recent work has suggested that the statistics of natural scenes differ between the upper and lower visual fields in terms of their distributions of luminance and luminance contrast. Using a natural scene database captured to reflect the visual environment of mice, Abballe and Asari (2022) found that the relative contrast in the visible and UV parts of the spectrum of natural scenes differ between the upper and lower visual fields, with the difference or “chromatic contrast” being larger in the upper than lower visual fields. Using signed contrast, Qiu et al. (2021) found that natural scenes were biased towards dark contrasts, especially in the upper visual field, and that this was paralleled by more light-offset-sensitive ganglion cells in the ventral mouse retina. A dark bias in natural scenes has also been reported in previous work on full-field natural images (Cooper & Norcia, 2015; Ratliff et al., 2010) and several physiological studies have found a dark bias in responses at the level of V1 (Jin et al., 2008; Xing et al., 2010; Yeh et al., 2009), particularly at low spatial frequencies (Jansen et al., 2019) and in the human visual evoked potential (Norcia et al., 2020). The work by Norcia et al. also demonstrated that an OFF-bias is present in both the UVF and the LVF, but this was only measured at a single suprathreshold contrast. Importantly, Qiu et al.’s data in mice suggests that the natural scene dark-bias may be field dependent, warranting an investigation of the ON/OFF biases in the UVF and LVF in human observers.

A necessary step towards understanding whether the visual system is adapted to prevailing scene statistics is the measurement of visual responses as a function of stimulus contrast. Laughlin (1981) showed that contrast responses of fly large monopolar cells could be directly related to the cumulative probability distribution of scene contrasts. This suggested that contrast coding in the fly efficiently mapped visual responses onto the distribution of contrast in natural scenes. Only three small-scale studies have measured contrast response functions for contrast increments and decrements (Kremkow et al., 2014; Rahimi-Nasrabadi et al., 2021; Zemon & Gordon, 2006), but they did not assess visual field asymmetry. The studies of Kremkow et al. and Rahimi-Nasrebadi et al. both report that OFF-contrasts were represented by a more linear contrast response function that failed to saturate, while the responses to ON-contrasts followed an accelarating and saturating non-linearity. Conversely, Zemon and colleagues reported the opposite: repsonses to OFF contrasts did saturate, and did so at a lower contrast than ON. Given the initial evidence that CRF shape may differ for increments and decrements, we asked whether these differences may also depend on visual field location. We find that ON and OFF contrast responses are highly nonlinear in the lower visual field, classically accelerating across low contrasts as contrast increases and saturating at high contrast. In the upper visual field, however, decremental/OFF responses are quasi-linear, while incremental/ON responses remain sigmoidal.

## Materials and methods

### Participants

Twenty-seven participants were recruited from the Stanford University Psychology department course credit pool (mean age = 23 yrs, SD = 5 yrs; 19 female). Participants were instructed to wear their most recent optical correction. Visual acuity was measured using a Bailey-Lovey chart (chart #5, Precision Vision, Woodstock, IL, USA) at 4m, and participants had to achieve a visual acuity of at least 0.2 LogMAR in each eye. Near-field stereo-acuity was assessed with a Randot Stereotest (Stereo Optical Co., Inc., Chicago, IL, USA) at 41cm with a passing score of 70 arc-seconds or better. One subject was assessed for Stereo-acuity using the Frisby stereo-test. Two near-sighted participants had forgotten their distance correction and failed the Bailey-Lovey chart at 4m, but were allowed to participate on the basis of passing the near-field stereo-acuity test at 41cm. Informed written and verbal consent was obtained from all participants prior to participation under a protocol approved by the Institutional Review Board of Stanford University.

### Visual stimuli

Contrast response functions for ON- and OFF-pathway biasing stimuli were measured for each observer in response to a hexagonal array of flickering probes. The entire array subtended 42°× 25° of visual angle (see Figure 1A). Two types of hexagonal elements are present in the array: probes and pedestals. Pedestal hexagons are larger elements that have a fixed luminance of 92.4 cd/m^2^, while probe hexagons are smaller elements within pedestals with luminance modulation that was experimentally manipulated (see the hexagonal elements in Figure 1B). Across 10 conditions, probe elements were temporally modulated (in achromatic luminance) according to a saw-tooth profile, the fast-phase of which was set to bias evoked responses either towards the ON or OFF pathway (Kremers et al., 1993). ON-pathway biasing stimuli were defined as probes that rapidly increased in luminance and slowly decreased, while OFF pathway biasing stimuli were defined as probes that rapidly decreased in luminance and slowly increased (see wave-forms in Figure 1B). All hexagons were presented against a low-luminance background of 15.2 cd/m^2^. All elements were scaled with eccentricity as detailed previously (Norcia et al., 2020).

**Figure 1:**
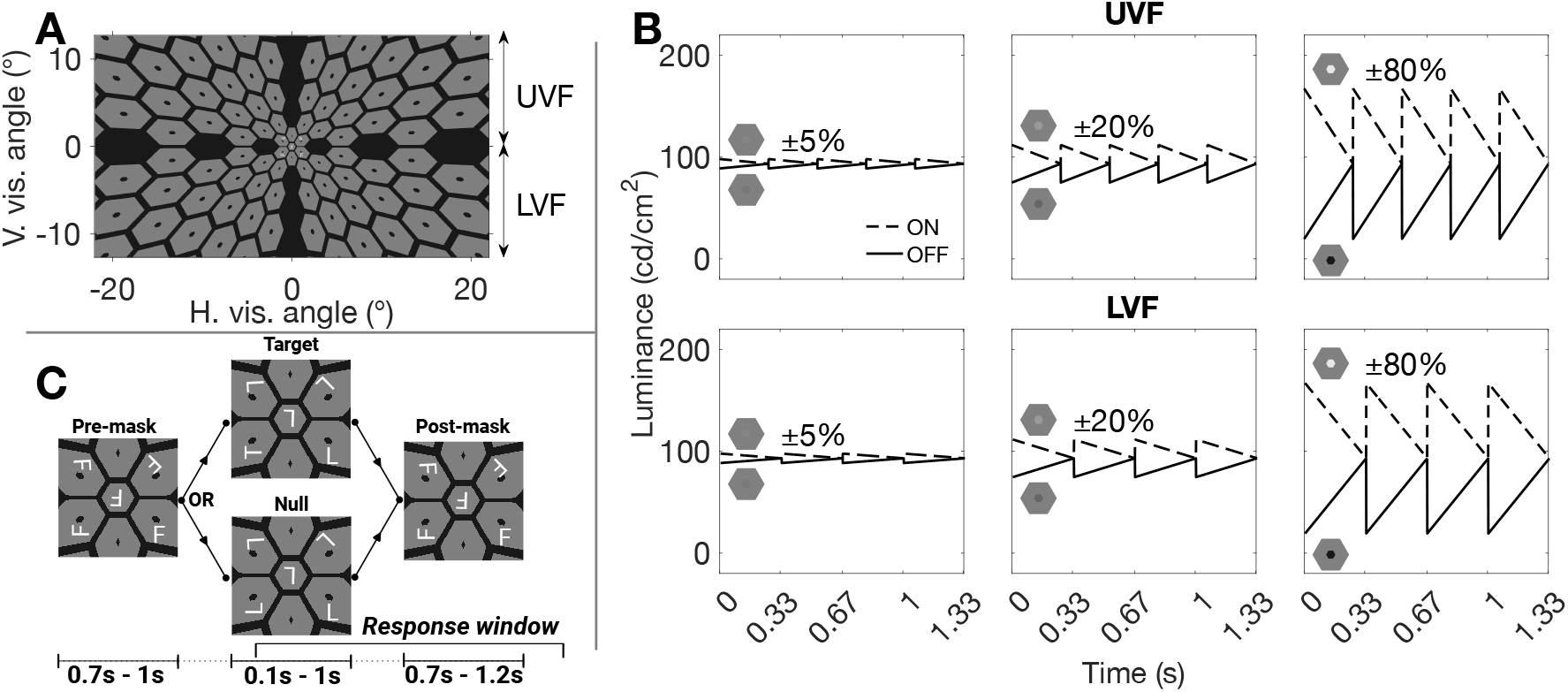
A single frame of the hexagonal stimulus array at 80% OFF contrast (A). The luminance of the central probes is varied to bias responses to the ON or OFF pathways at different Weber contrasts. The temporal frequency of the saw-tooth stimulus was different for the UVF and LVF, which allows for the spectral decomposition of these signals (B). A schematic overview of a single trial of the concurrent attention task (C).

In order to record Steady-State Visually Evoked Potentials (SSVEPs) simultaneously from the UVF and LVF, probes in the UVF flickered at 3.75 Hz, and probes in the LVF at 3 Hz (see right-most ordinate of 1A). All probes in a given half-field were modulated synchronously with an identical temporal waveform. We chose 3Hz and 3.75Hz for several reasons: First, these frequencies are close to each-other, such that human temporal contrast sensitivity is similar for the two stimuli. Second, they can be easily rendered on a 60Hz display (3Hz is exactly 20 frames, 3.75Hz is 16 frames). Third, they have a high least common multiple (15Hz), so harmonic frequencies can be independently interpreted up to 15Hz. Finally, low frequencies (<5Hz) allow the main temporal components of the response to evolve and mostly return to baseline over a single stimulation cycle. Offline spectral decomposition was used to separate responses at these frequencies (and their harmonics), effectively obtaining independent responses for the simultaneously presented UVF and LVF probes. SSVEPs were measured at 5 geometrically spaced contrast levels (5, 10, 20, 40, & 80%), with contrast defined using the Weber definition: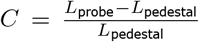, where *L* is the luminance of the probe or the pedestal. There were 10 stimulation conditions (2 pathways × 5 contrasts), with 9 trials per condition (90 trials total). A single trial consisted of 16 seconds of continuous stimulation for a given condition, with a 2.5s - 3.5s background-only period in-between trials. Longer breaks, typically lasting approximately 2 minutes, were provided every 30 trials. Three participants only completed 6 trials per condition (60 trials total). To control participant vigilance and clamp attention at a more constant value, a letter task was presented concurrently during steady-state stimulation. Five letters were presented within the central 2 degrees of the display, one central letter flanked by 4 letters (see Figures 1A and 1C). Each letter element subtended approximately 30 minutes of arc. In a single trial of the task (lasting up to 3.2s) there were three phases (see Figure 1C), the pre-probe mask, the probe, and the post-probe mask. The pre- and post-probe mask phases were identical arrays that contained only “F”s, while the probe phase could be a “null” that contained only “L”s, or a “target” that contained four “L”s and one “T”. The participant was instructed to respond with a button press when they saw the target. The position of the letter “T” was randomised across trials. The duration of the target probe was titrated with a two-down-one-up staircase, such that the task quickly converged on a threshold (minimum presentation time was 100 ms, the max was 1 s). The duration of the pre- and post-probe mask was randomized on every trial. The pre-probe mask lasted between 0.7 and 1 seconds, and the post-probe mask lasted between 0.7 and 1.2s. The orientation of the letters was randomized on every trial of the attention task. Within a trial, the letter orientations were held constant at each letter location. Trials were presented repeatedly for the duration of the flickering stimulus, and the value of the staircase was carried across the trials of the flickering stimulus.

### EEG recording

The EEG was recorded using a 128-channel EGI HydroCel SensorNet and NetStation 5.2 software at a sample frequency of 500 Hz and resampled to 420 Hz (7 samples per video frame). Every effort was made to reduce channel impedance below 100 kΩ at the point of data collection, and impedance was checked in-between trial blocks, with electrodes rewetted if necessary.

### Artifact rejection and EEG filtering

For each observer, the raw EEG was amplified (gain = 1000, 24-bit resolution) and digitally filtered with a 0.3 – 50 Hz band-pass filter. The data were then artifact rejected with in-house software written in Objective C using the following criteria. First, consistently noisy individual channels were detected, rejected, and substituted with the average of the six nearest-neighbour channels. Channels were classified as consistently noisy if over 15% of samples exceeded 30 µV (excluding breaks). After this, the data were re-referenced to the common average. Second, the 16-second trials were broken down into the maximum number of “subtrials” that still possessed an integer number of cycles at both stimulus frequencies. For each trial, this yielded 12 sub-trials of 1/.3 seconds (4 cycles of 3 Hz and 5 cycles of 3.75 Hz). The first and last sub-trial of each trial were always discarded. Third, to reject data containing coordinated muscle movements and blinks, 1/.3 second-long sub-trials were excluded for all channels if more than 5% of channels exceeded an amplitude threshold of 60 µV. Fourth, 1/.3 second sub-trials of individual channels were excluded if more than 10% of samples exceeded 30 µV. These light-touch artefact rejection criteria were derived empirically for adults over hundreds of previous recordings. They readily pick out muscle and blink artifacts as well as electrode motion artifacts which are not of neural origin, leaving relatively clean EEG. Finally, any subjects who had more than 30% of their samples rejected (after channel substitution) were entirely removed from the analysis. Six subjects failed this final criteria, meaning 21 were used in the forthcoming analysis.

### Spectral Analysis

Spectral analysis was performed for each participant, for every sensor, at the sub-trial level using a recursive least squares (RLS) filter (Tang & Norcia, 1995). Briefly, RLS is equivalent to the discrete Fourier transform (DFT) but is more effective when short trial lengths are used. Conceptually, RLS directly fits sine and cosine waves at selected stimulus-relevant frequencies to a time-series. This process yields complex-valued estimates of harmonic amplitudes that can be used in the same way as the amplitudes from the DFT. In the present work, RLS was performed up to the 3rd harmonic for each stimulus frequency. Assuming a participant had no sub-trials rejected, this yielded 90 spectral estimates per condition, visual field location, harmonic, and participant (60 in the participants who completed 6 trials of each condition). It should be noted that the multi-frequency stimulation we have employed makes the display of conventional VEPs more challenging, as the resultant wave-forms are a complex mixture of the two stimulation frequencies. Spectral analysis can “un-mix” these signals, meaning the frequency domain representation of the data is more conducive to interpretation. See Figure 2, where we have plot both the frequency and time domain representation of the cross-participant average response to an 80% OFF-contrast flicker at channel 75. Note, that we use an “xFy” nomenclature, such that “1F2” refers to the 1st harmonic of the 2nd stimulus frequency (3Hz). The LVF and UVF VEP is not readily discernible from the time-domain representation of the data (right-most panel), but the frequency domain representation readily separates LVF and UVF spectral peaks at 3Hz and 3.75Hz, respectfully (and their integer multiple harmonics).

**Figure 2:**
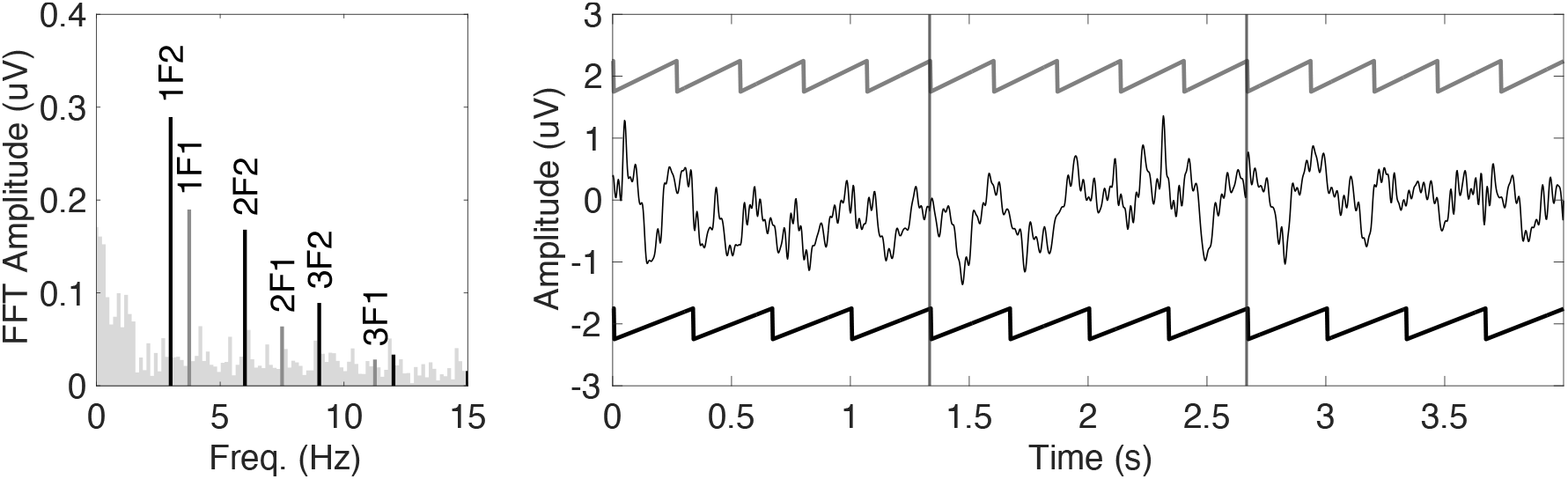
Frequency domain (left panel) representation of the average (N=21) response to an 80% OFF-contrast flicker at channel 75 (approx. occiptial pole) for 6.6s of data. The stimulus related frequencies are labelled up to the 3rd harmonic. The right panel is the time-domain representation of the same data, but the x-axis has been limited to 4s to aid visualisation. Each vertical reference line shows the time at which an integer number of stimulus cycles were completed for both stimulation frequencies (see the reference saw-teeth at 3Hz (LVF) and 3.75Hz (UVF)).

### Normalisation and dimension reduction

Despite the use of identical stimuli, the amplitude of the SSVEP varies between individuals. This may be partially driven by differences in skull and scalp thickness/conductivity. These passive electrical differences would not only scale the stimulus evoked response, but also the associated intracranial EEG noise. Therefore, in an effort to at least partially account for cross-subject differences in passive electrical properties, we normalised participants’ RLS complex-valued spectral data by an estimate of their noise level. To ensure we did not introduce any condition-wise bias, each subject’s spectral data was scaled by a single unique value derived from the entirety of their data. This value was calculated as follows: first, for the real and imaginary part of every stimulus harmonic, we calculated the variance across all sub-trials. Then, we took the mean of the real and imaginary variance (this is equivalent to Eqn. 2 of Victor & Mast (1991)), and took the mean of this value across harmonics. See Equation 1, below:

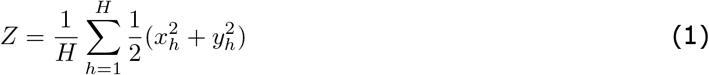

Where *H* is the number of harmonics, and 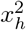 and 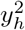 are the variances of the real and imaginary parts for the *h*^*th*^ harmonic. This was done separately for each condition, such that *Z*_*c*_ represents the total variance of condition *c*. As shown in Equation 2, we averaged this value across conditions and took the square root to return to units of microvolts. This process was repeated for all subjects, such that *M*_*n*_ represents the normalizing denominator *M* for the *n*^*th*^ subject. Finally, for each subject, we divided the raw real and imaginary components of all trials in all conditions by this value. This process leaves us with normalized 128-channel spectral data for each participant. Effectively, we have ‘z-scored’ the RLS spectral estimates.

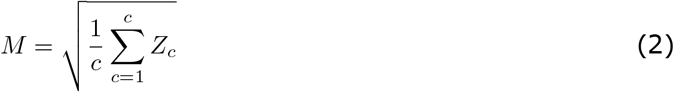

After normalisation, Reliable components analysis (RCA) in the frequency domain was used to reduce the dimensionality of the 128-channel data to a smaller number of more easily interpreted components, as previously detailed (Dmochowski et al., 2015). Briefly, each reliable component (RC) is a weighted sum of electrical potentials across all channels. The weight vectors are derived through an eigenvalue decomposition performed on a 128 × 128 matrix where each element represents the ratio of within-trial covariance (*R*_*xx*_) to cross-trial covariance (*R*_*xy*_). Solving this decomposition provides multiple ranked components (i.e. spatial filters) that maximise *R*_*xx*_/*R*_*xy*_, with the 1st RC containing the maximal contribution from channels with consistent cross-trial activity. This reflects a fundamental quality of the SSVEP, where repeated presentations of the same stimulus produce similar stimulus-locked neural activation across multiple trials. For the present analysis, RCA filters were trained at the group level on the normalized RLS data for 80% contrast, but separately for the frequencies related to the UVF and LVF (up to the 3rd harmonic). This yielded a set of spatial filters for both the UVF and the LVF through which the normalized RLS estimates of all conditions were then projected. We only analyse the first three RCs for each visual field location. After projection through a given RC filter, at the individual subject level, we took the vector mean for all harmonics and all conditions across sub-trials, took the absolute value, and calculated the root-mean-square (RMS) amplitude across harmonics. This leaves us with a single unsigned amplitude value for every condition and visual field location, for three RC topographies, for every participant. Importantly, these values are compatible with univariate model fitting/statistics.

### Model fitting and statistics

To provide a compact description of ON and OFF pathway responses from the UVF and LVF, the group-level mean of observers’ RMS contrast responses (post-RCA projection) was fitted with a hyperbolic ratio function (Equation 3), which is often used to model nonlinear responses to contrast (Albrecht & Hamilton, 1982). This function has the useful quality of describing a range of response profiles with only four parameters: *rMax, c*50, *n*, and *rMin*. The *rMax* is a scaling coefficient that describes the theoretical saturating response of the responsive neural population. All else being equal, increasing *rMax* is akin to increasing “response gain” - stretching the function along the output axis, such that the same range of input values are encoded by a greater range of outputs (increasing response granularity). Broadly, the *c*50 describes the contrast at which the neural population reaches half of the value of *rMax*. Increasing the *c*50 reduces input gain, stretching the response function to encode a wider range of contrasts with the same range of outputs (reducing response granularity). The exponent *n* describes the form of the non-linearity occurring proximal to the *c*50. Exponents greater than 1 describe the classic accelerating-then-saturating non-linearity, while exponents of 1 or less describe a purely saturating function. Finally, the *rMin* is an additive constant that, in the case of EEG, simply describes the noise floor.

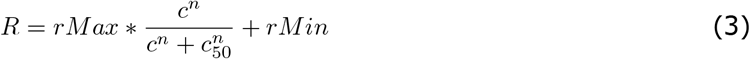

To enable statistical comparisons of the fitted group-level parameters between conditions, bootstrapped confidence intervals were generated on the fits. For each condition, we did the following using a participant (*n*) by contrast (*c*) matrix of RMS amplitudes. In a single iteration, the rows of *n* were re-sampled with replacement and the mean taken along the *n* dimension. We then fit the model (Eqn.3) to this 1 x *c* vector of amplitudes and saved the parameters. This was repeated 20,000 times. For each condition, the re-sampling seed was reset to the same value, such that the participants selected were the same in the *x*th draw of any condition. To obtain 95% confidence intervals (CIs) on the fit parameters, we took the 2.5th – 97.5th percentiles of the parameter distributions. To obtain 95% CIs on the differences between conditions, we took the same percentiles on the differences between each of the 20,000 rows for two given conditions. Where these CIs do not contain zero, a significant difference between two conditions can be concluded. To draw shaded confidence regions around the fit to the empirical mean, for each bootstrap draw, we evaluated the hyperbolic ratio fit at discrete values of contrast (1% increments). The 68% CI of these pseudo curves were then refit with Equation 3 and plotted as 1 standard-error bounds of the fit around the empirical mean. We also performed an ANOVA as an additional description of effects. A semi-parametric 3-way (contrast x pathway x VF) repeated measures ANOVA was carried out on RMS amplitudes using the RM() command of the MANOVA.RM package (Friedrich et al., 2019) in the R programming language. For repeated measures designs, this package provides a permutation approach for calculating “Wald-type statistic” (WTS) p-values. The permuted WTS is robust to violations of normality and performs well at low-to-moderate sample sizes (Friedrich et al., 2017). Post-hoc ANOVA tests were corrected for multiple comparisons using the Bonferroni-Holm method (Holm, 1979). While this ANOVA cannot speak directly to differences in nonlinear response properties, it can reveal gross level differences in the data (i.e. are responses to OFF-biasing stimuli generally larger than responses to ON-biasing stimuli).

## Results

### Response topographies are consistent with retinotopic organisation

The frequency tagging approach will produce separable responses for the UVF and LVF in retinotopic areas of the brain. In Figure 3, we show the grand-average RLS spectra of the channel showing peak amplitude for the UVF (left-hand panel) and the LVF (right-hand panel). These channels were selected by searching for the maximum cross-subject vector mean RLS amplitude at each fundamental frequency used in this experiment (1F1 and 1F2) at 80% contrast for OFF stimulation. There are two inlaid axes in each panel. The left inlay shows a zoomed-in high-resolution spectrum calculated using the fast Fourier transform (FFT). The right inlay shows the topography of RLS responses at the first harmonic of the relevant stimulation frequency (again, at 80% OFF). Clearly, for the LVF, there is peak signal over more posterior channels, while the UFV representation is shifted towards more anterior/dorsal channels. This topography is consistent with responses that originate from retinotopic brain regions, and broadly matches the topography predicted by the forward modelling work of Ales, Yates, & Norcia (2010). In the RLS Spectra of Figure 3 there are well-defined peaks at stimulus-relevant frequencies (black and mid-grey bars), while the non-stimulus frequencies (lighter bars) are relatively flat. The abutting non-stimulus frequency bins in the inlaid high-resolution spectra also show no evidence of signal leakage. These spectra and topographies indicate that the timing of our stimulus was well-calibrated, and assure us that we are obtaining separable and topographically distinct responses from the UVF and LFV. That the UVF and LVF responses are independent is also supported by the absence of signal at inter-modulation frequencies. This also indicates that contributions to the VEP from wide-field receptive fields that are subtended by multiple probes is minimal.

**Figure 3:**
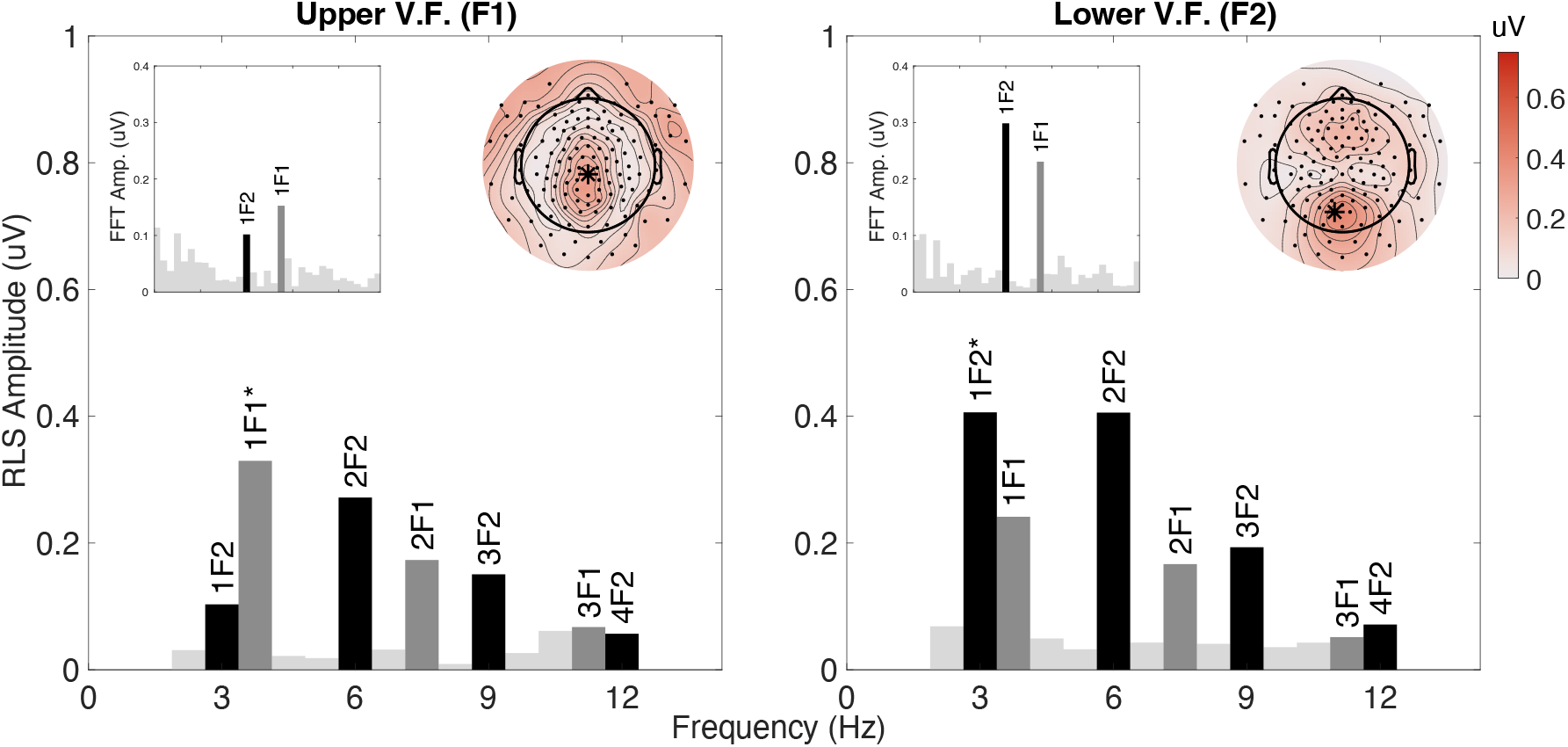
Grand-average (N=21) RLS spectra for two channels with maximum RLS-amplitude at fundamental stimulation frequencies. Data shown are responses to 80% contrast OFF-biasing modulation. The spectrum in the left panel is for channel 55 (max. for 1F1/UVF), while the right panel is for channel 71 (max, for 1F2/LVF). Bars colored grey and black highlight the frequencies related to the UVF and LVF, respectively. The lighter-gray bars are non-stimulus frequencies. The inlaid axes of each panel show higher-resolution FFT spectra (left inlay) and RLS topographies of the fundamental frequencies used in the experiment with the peak sensor highlighted by an asterisk (as is the generative spectral peak) (right inlay)

To reduce these multi-channel data to analytically tractable components, we used RCA to reduce the channel dimension to a smaller set of reliable components (RCs). In Figure 4 a scree plot illustrating the proportion of cross-trial covariance accounted for by the first 12 RCs recovered for the upper and lower visual fields is presented. The first RC for both field locations captures more than twice the covariance than the next reliable component. The forward model topographies for the first three RCs are shown inlaid in this figure. Note the similarity between the group-average topographies presented in Figure 3 and their RC1 counterparts presented in Figure 4.

**Figure 4:**
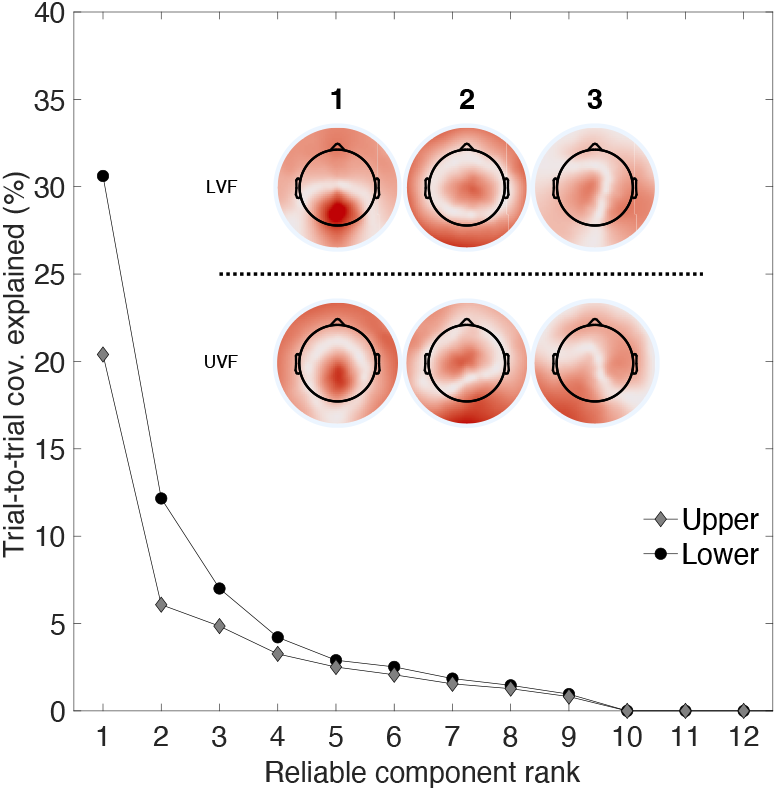
Scree plot showing trial-to-trial covariance explained by the first 12 reliable components for the upper (diamonds) and lower (circles) visual fields. Inlaid topographies show the forward model projections of the first 3 components, plotted on the same color-scale (the units are arbitrary).

### Absence of OFF/decrement saturation in the UVF

We first focus on the difference between ON- and OFF-pathway biasing responses separately for each visual field location. In Figure 5 we show the contrast response functions derived from RC1 for the upper (left panel) and lower (right panel) visual fields. In both locations, the OFF pathway amplitudes trend higher than the ON pathway amplitudes at all but the lowest contrast, especially in the LVF. Additionally, the shape of the fitted response functions (solid curves) indicates that the LVF responses - both ON and OFF - show an accelerating and saturating non-linearity, but with OFF-pathway responses growing at a greater rate with respect to contrast. In the UVF, the ON-pathway shows a similar sigmoidal non-linearity, but the OFF pathway shows little evidence of saturation. Indeed, the responses from both pathways in the UVF are quite similar at low and intermediate contrasts.

**Figure 5:**
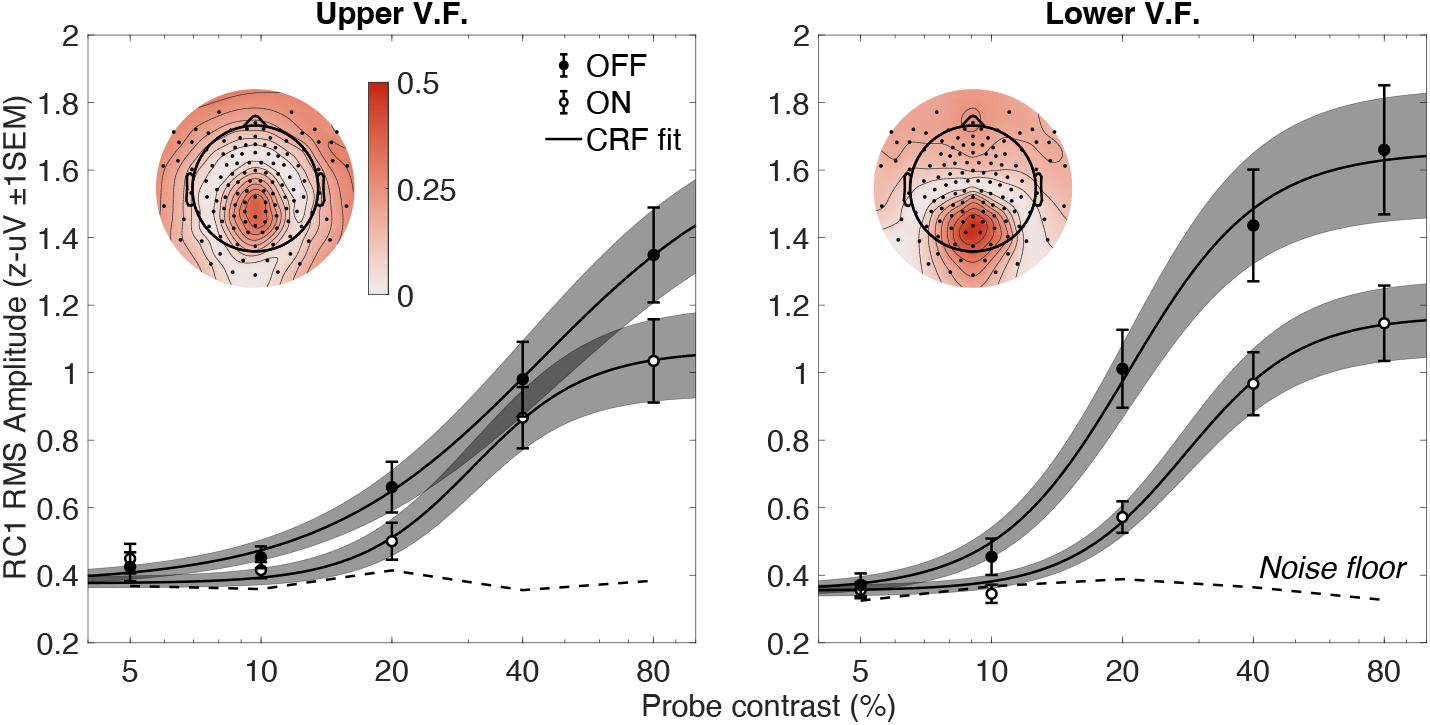
Group-level (N = 21) RC1 Topographies and contrast response functions. Shaded regions illustrate 68% confidence intervals on the model fit to the empirical mean. Error bars are 68% confidence intervals on the empirical mean response for a given contrast. The units on the color-bar are micro-volts.

To test the statistical significance of these response patterns, we first conducted a semi-parametric 3-way repeated measures ANOVA on the RMS values contributing to the means shown in Figure 5. This analysis reported significant main effects of contrast (WTS(4) = 60.80, *p* < .001) and probe polarity (WTS(1) = 38.56, *p* < .001), but a non-significant main effect of visual field (WTS(1) = 3.35, *p* = .084). There was significant 3-way interaction between contrast, polarity, and visual field (WTS(4) = 14.65, *p* < .05). To elucidate this 3-way interaction, we conducted additional 2-way repeated-measures ANOVAs, first for visual field location. In the UVF, there was a significant effect of contrast (WTS(4) = 42.43, *p* < .001) and polarity (WTS(1) = 21.957, *p* < .001) on response amplitude. The same was true in the LVF for contrast (WTS(4) = 51.772, *p* < .001) and polarity (WTS(1) = 28.1, *p* < .001). There was a significant interaction between contrast magnitude and contrast polarity in the UVF (WTS(4) = 20.23, *p* < .05) and the LVF (WTS(4) = 23.57,*p* < .05). This result supports our observation that the OFF response is generally larger than the ON response, but that the difference between the ON and OFF pathways depends on the contrast being tested in both visual field locations. Next, we marginalised contrast polarity. In both pathways, the main-effect of contrast magnitude was significant (ON: WTS(4) = 54.94, *p* < .001; OFF: WTS(4) = 59.62, *p* < .001). In the ON pathway, the effect of visual field was non-significant (WTS(1) = 0.26, *p* = .615), but it was significant in the OFF pathway (WTS(1) = 5.81, *p* < .05). In both pathways, the interaction between visual field location and contrast magnitude was non-significant after correction for multiple comparisons. This analysis suggests that OFF-pathway responses are generally larger in the LVF than in the UVF, and that the ON-pathway response is more similar between the UVF and LVF. However, this analysis cannot easily describe differences in the shape of contrast responses. For this purpose, we proceed to the analysis of fitted contrast response functions.

To aid in the interpretation of parameter fits, the fit parameters to the empirical mean responses and parameter histograms (from the bootstrapping procedure) are presented in Figure 6. As stated in the the methods section, a significant difference is concluded when the bootstrapped confidence interval on the difference between two conditions does not contain zero. These difference histograms are shown in Supplementary Figure 9. Visual assessment of Figure 5 indicates that the response functions of the ON and OFF pathways in the LVF are similar, but the OFF response is stretched along the output axis (elevated rMax) and shifted leftwards on the input axis (reduced c50). That is, the curves differ in scale and location, but both accelerate through lower contrasts and saturate at higher contrasts. Analysis of the parameter fits supports this observation - in the LVF, the OFF pathway shows a significantly lower c50 (*p* <.05) and a significantly higher rMax (*p* <.01), but the distributions of exponents practically overlay between the ON and OFF-pathways. This indicates that both pathways show a similar form of non-linearity, but the range of contrasts over which the curves are most sensitive (the c50) and the degree of sensitivity (the rMax) differs.

**Figure 6:**
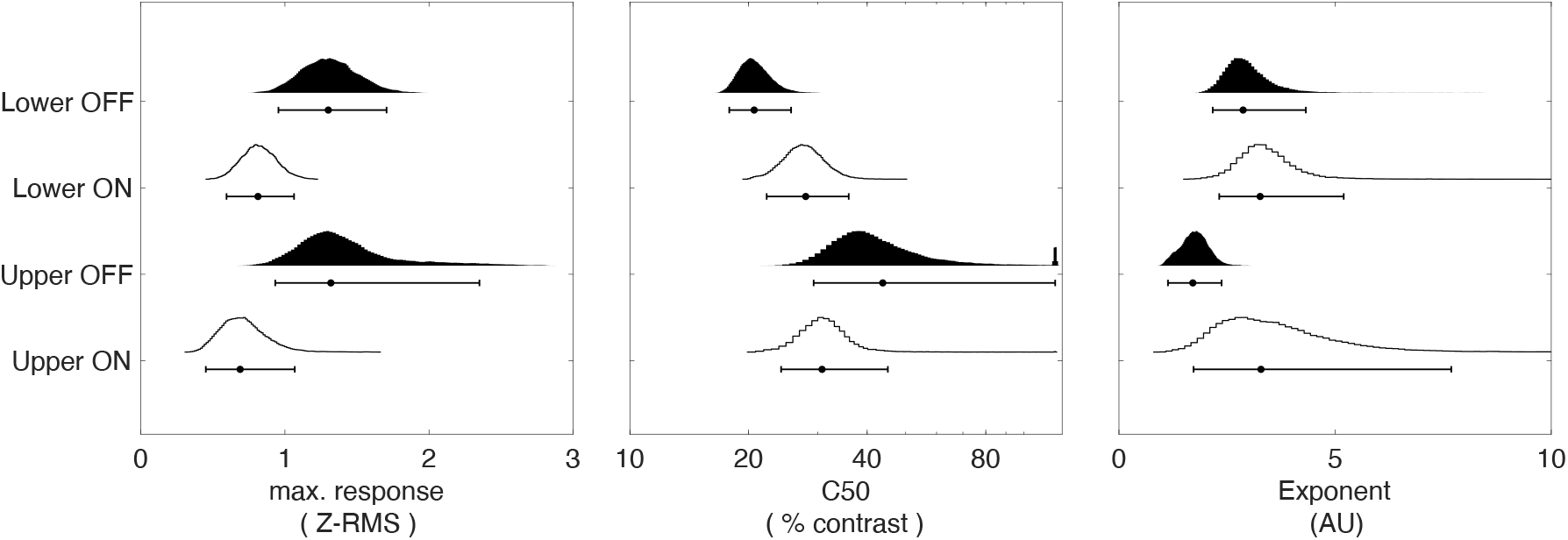
Half-violin plots showing parameter estimates produced by the group level fit bootstrapping procedure for RC1. Histograms have been color-coded by the pathway being biased (White = ON, Black = OFF). The parameter displayed in each sub-plot is shown in the x-axis labels.

In the UVF, the OFF-pathway fits shows little evidence of saturation, while the ON pathway fit saturates much like it does in the LVF. Indeed, between the LVF and UVF, there are no significant differences between any ON-pathway fit parameters. In the UVF, the OFF pathway c50 distribution has a long tail towards higher contrasts, with approximately 32% of c50s in excess of 50% contrast, and 7% reporting c50s beyond 100% contrast. In all other conditions, no more than 1% of c50s are beyond 50% contrast. It should also be noted that the exponent value in the UVF-OFF is significantly lower (*p* < .05) than that of both LVF pathways. When coupled with its low exponent value, that the OFF-UVF response does not reach half its maximum by 50% contrast indicates that a significant proportion of the fitted curve’s dynamic range is outside of the range of physically possible OFF contrasts. That is, the probe elements being displayed could not possibly get any darker in the limit of ambient light incident upon the display. This suggests that the OFF pathway in the UVF encodes contrast very differently than in the LVF: rather than having a narrow dynamic range with high sensitivity, it has a broad dynamic range of lower sensitivity. Importantly, with c50s so high, the interpretability of the rMax and exponent parameters is diminished. This is because the model we have fit is saturating at a minimum, and accelerating and saturating depending on the model parameters. Here, the OFF-pathway in the UVF has a significantly higher rMax than the ON-pathway (*p* < .05), but is is clear that is is from a failure to saturate, rather than a re-scaling of the same response curve (as in the LVF). Overall, the ON and OFF pathways are clearly distinguishable in the Lower visual field: they have a similar function shape, but differ in range of contrasts over which they are most sensitive. In the Upper visual field, our results are more complex. Here, the ON and OFF pathways are almost overlapping at low to moderate contrasts, but subtly diverge at higher contrasts, seemingly failing to saturate in the OFF pathway.

It is possible that these results are specific to RC1. Reliable components 2 and 3 have different topographies that may reflect different neural generators with different tuning properties. To examine this possibility, we analysed the contrast responses obtained when the data are instead projected through RC2 (Figure 7), First, note the reduction in the overall amplitude of these responses - approximately 50%. Because the noise-floor remains the same regardless of the visually evoked signal, these smaller responses are more difficult to compare. Nevertheless, the coarse pattern of the results present in RC1 are maintained. Responses grow with increasing contrast, the LVF response appears larger than the UVF response, and within each visual field location, the OFF-biasing response appears to be larger than the ON-biasing response. This is supported by another 3-way repeated measures ANOVA, that shows a significant main effects of contrast ((WTS(4) = 31.11, *p* < .001), visual field (WTS(1) = 10.02, *p* < .01), and a significant main effect of contrast polarity (WTS(1) = 18.35, *p* < .001). Unlike in RC1, however, there are no significant 2- or 3-way interactions between contrast, probe polarity, and visual field location. Additionally, the confidence intervals on the model fits entirely overlap in RC2, suggesting that the responses are quite variable across participants. Indeed, the c50 and exponent fits show more variability than those of RC1 (see Figure 8 - note the changed abscissa limit for the exponent). Only the difference in rMax between the ON and OFF pathways in the LVF is significant (the difference distributions are shown in supplementary Figure 2). It is likely that RC2 represents some small contribution from a later source in the visual processing stream, but we do not possess a sample sufficient to fully characterise it. This issue is compounded further in RC3, where the model fit distributions become bimodal (data not shown). For this reason, we will focus on the discussion RC1 for the remainder of this work, as it demonstrably describes most of the stimulus-locked activity present in our data.

**Figure 7:**
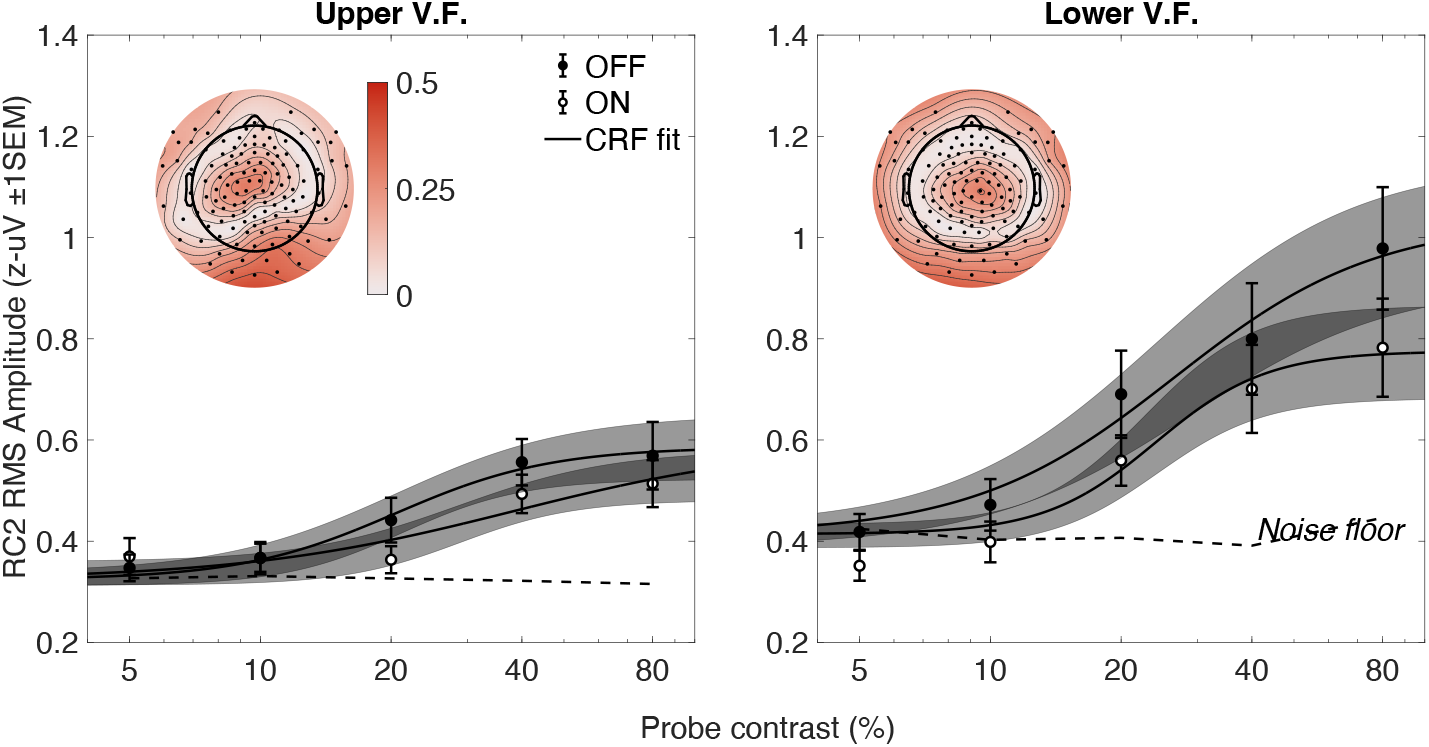
Group-level (N = 21) RC2 Topographies and contrast response functions. Shaded regions illustrate 68% confidence intervals on the model fit to the empirical mean. Error bars are 68% confidence intervals on the empirical mean response for a given contrast. The units on the color-bar are micro-volts.

**Figure 8:**
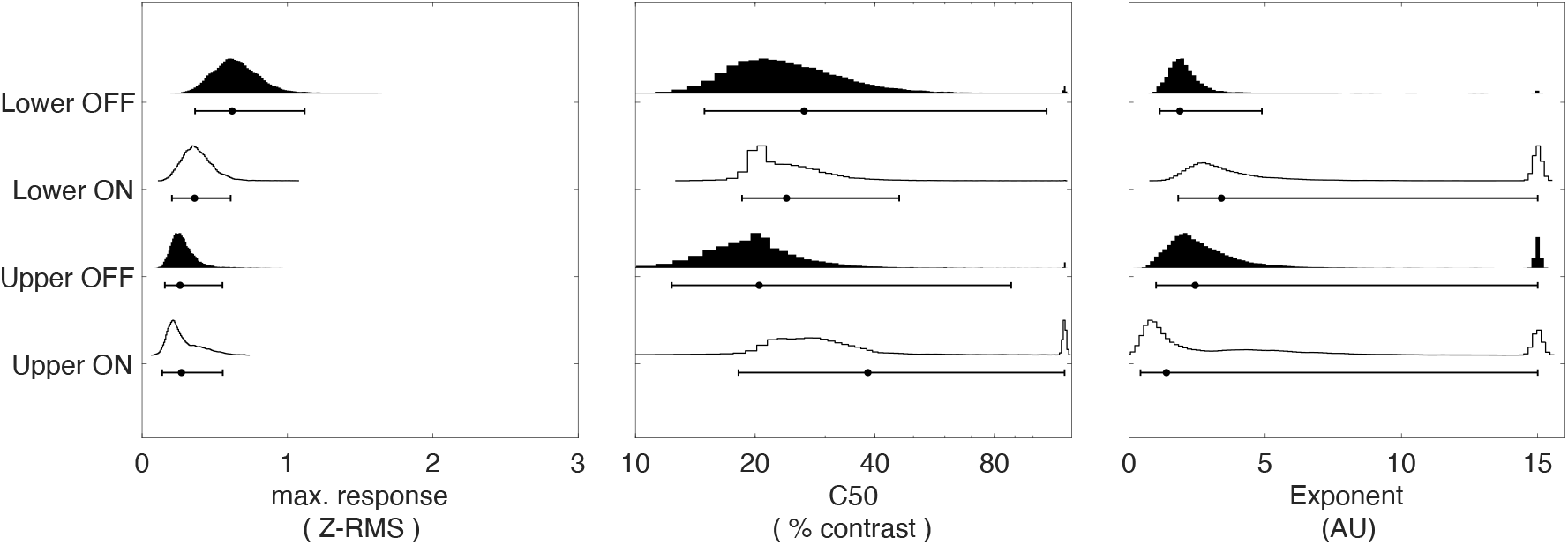
Half-violin plots showing parameter estimates produced by the group level fit bootstrapping procedure for RC2. Histograms have been color-coded by the pathway being biased (White = ON, Black = OFF). The parameter displayed in each sub-plot is shown in the x-axis labels.

## Discussion

In our previous work, we reported that the OFF pathway SSVEP is higher in amplitude than the ON pathway SSVEP in response to sawtooth stimuli in both the UVF and the LVF (Norcia et al., 2020). Here, we replicate this result, but demonstrate that the degree to which this is true depends on the stimulus contrast in conjunction with the visual field location being tested. In the LVF, we find that both the ON and OFF pathway-biased responses show clear sigmoidal non-linearities, with the OFF pathway response being larger than the ON pathway response and saturating earlier. In the UVF, only the ON pathway shows a reliable sigmoidal non-linearity, while the OFF pathway encodes contrast quasi-linearly, showing little saturation behaviour. These results suggest that full-field assessment of ON and OFF pathway function provides an insufficient description of the sensitivity profiles of these pathways, and encourages a further exploration of why these pathways may have a visual field dependence.

### Comparison with human ON/OFF contrast response functions

Although several reports have measured VEPs in humans using ON- & OFF-pathway biasing stimuli (Mutlukan et al., 1992; Roveri et al., 1997; Zemon et al., 1995; Zemon et al., 1988), very few have measured contrast response functions (Kremkow et al., 2014; Rahimi-Nasrabadi et al., 2021; Zemon & Gordon, 2006), and none have investigated visual field asymmetry. Kremkow et al. (2014) and Rahimi-Nasrabadi et al. (2021) both measured CRFs using full-field chequerboards, biasing responses towards the ON or OFF pathway by fixing the luminance of background checks and varying the luminance of decremental/incremental target checks. Kremkow and colleagues used stimuli with binary backgrounds (black or white), while Rahimi-Nasrebadi et al. used mean-luminance backgrounds. Despite differing stimulus parameters, their results are in broad agreement: responses to incremental stimuli showed the strongest saturating non-linearity, while the decremental responses failed to saturate. To first order, their results imply that the OFF pathway is optimised to encode a broad range of contrasts with moderate accuracy, while the ON pathway encodes a discrete range of contrasts with high accuracy. While this is consistent with our UVF results, it is not consistent with our findings in the LVF, nor the full-field results of Zemon and Gordon (2006). Zemon and Gordon used a frequency domain approach, sinusoidally modulating the luminance of isolated checks on a mean-luminance background. Overall, the amplitude of the response at the fundamental frequency was highest for the OFF biasing stimuli, and the decrement responses saturated earlier than increment responses with both showing sigmoidal non-linearities. Taken together, our results from the LVF more closely match the pattern reported by Zemon and Gordon, while our results in the UVF are closer to the pattern reported by Kremkow et al. (2014) and Rahimi-Nasrabadi et al. (2021). It is difficult to find an explanation for the differences between these studies, as they used different stimuli that may bias responses towards neural populations with different tuning properties.

Another difference between ON and OFF pathways is their spatial tuning. Two reports have measured incremental & decremental contrast response functions for full-field stimuli, varying element and stimulus size (Kremkow et al., 2014; Zemon & Gordon, 2006). Kremkow and colleagues found that the OFF responses dominated in amplitude at lower grating spatial frequencies, and that the ON pathway responses had a higher preferred spatial frequency than OFF. This elevation of ON-pathway spatial frequency tuning was attributed to “neural blur” caused by an early transducer nonlinearity unique to the ON-pathway (we report a nonlinearity in both pathways depending on visual field location). Conversely, using isolated checks, Zemon and colleagues reported a broader spatial tuning for the OFF/decremental stimuli, with OFF-responses being higher for denser check grids. The results of these reports may seem at odds, but Zemon and colleagues findings may also be explained by ON-pathway neural blur, where more densely packed checks would be less distinct. At the very least, this demonstrates that the ON and OFF pathways are likely differentially tuned to the spatial properties of the perceived world. While possible interactions between size tuning and visual field location have not been investigated using incremental and decremental stimuli, the visual field dependence of contrast response function shape has been investigated, but using unsigned contrast. Laron and colleagues (2009) investigated the visual field asymmetry of contrast response functions as a function of eccentricity using the multi-focal VEP. They found that responses to foveal probes saturated very late (c50 >50% contrast), and that c50s reduced substantially with increasing eccentricity. While their data cannot speak to differences between the UVF and LVF (because they pooled over polar angle), it demonstrates that contrast response functions do vary by eccentricity, as we have shown for polar angle using signed contrast stimuli.

The diversity of stimulus conditions may have contributed to the discrepant results found in previous studies. The size of stimulus elements used varied widely between reports: Zemon and Gordon (2006) reports using 9 arcmin isolated checks, Rahimi-Nasrabadi et al. (2021) used 30’ checks. Kremkow et al. (2014) did not report the check size they used. Laron et al. (2009) used dartboard stimuli scaled for cortical magnification that are designed to equate the cortical representation of stimuli presented at different eccentricities, but the other studies did not. This may mean that the presented stimuli will have been optimal for different eccentricities. Finally, it should also be noted that these studies all suffer from a low sample size (N=3-6), so some variation may simply be driven by unrepresentative samples.

### On the use of rectified stimuli

It is possible that our results are specific to the rectified stimulus waveform we have chosen to use. Presently, the saw-tooth probe is always modulating above or below the pedestal luminance, such that the direction of the fast phase of the saw-tooth is always consistent with spatial center-surround contrast of the probe with respect to the pedestal. A side effect of this definition is that the temporal mean luminance of the probe is different for ON- and OFF-pathway biasing stimuli, and this difference increases with contrast. Alternatively, one could modulate the probe symmetrically around the luminance of the pedestal. This would equate the temporal mean luminance across all polarities and contrasts, but the centre-surround contrast would be ON-center for half of the time, and OFF-center for half of the time. Out of an abundance of caution to not mix ON and OFF-pathway responses, we chose the rectified definition for the present work.

To what extent could the different ON- and OFF-pathway responses we have found be due to our choice of stimulus? There is a small corpus of literature on this matter. Prior psychophysical work has indicated that flicker detection thresholds are elevated when using rectified wave-forms (with higher temporal-mean luminance) instead of symmetric wave-forms (Anderson & Vingrys, 2000; Zele & Vingrys, 2007). This could mean that the smaller response of the ON-pathway we find may be due to the ON-biasing stimuli having higher temporal mean luminance than the OFF-biasing stimuli. Indeed, at a single contrast, we have previously shown that the use of a symmetric saw-tooth wave-form can reduce the difference between the VEPs elicited by the ON and OFF pathways, but we also demonstrated that this effect depends on the overall luminance of the display (Norcia et al., 2020). At a spatial mean luminance of 40 cd/m^2^, the OFF-pathway response was larger and faster for both symmetric and rectified wave-forms, challenging the notion that the difference is purely due to a difference in temporal mean luminance. However, at a higher mean luminance of 90 cd/m^2^, the amplitude and speed advantage for OFF-biasing stimuli was severely reduced for symmetric waveforms only. This result suggests that this is not simply a case of rectified vs symmetric stimuli, but rather that some luminance information is retained and used to alter ON- and OFF-pathway symmetry. Modelling and empirically mapping this parameter space is a worthwhile pursuit, but beyond the scope of the present work. Presently, the degree to which our finding generalises to alternative saw-tooth definitions and spatial mean luminance values is unknown.

### Relating CRFs to natural scenes

Information theory suggests that differences in ON/OFF pathway spatial tuning may be driven by the distribution of ON & OFF information in natural scenes (Laughlin, 1981). Indeed, there is evidence for differential ON/OFF distributions at the full-field level. By convolving a centre-surround receptive field model of variable size with natural scenes, Ratliff et al. (2010) demonstrated that OFF contrasts are significantly more common at all spatial scales. Using a similar approach, but varying receptive field properties, Cooper and Norcia (2015) demonstrated that OFF contrasts dominate particularly at low spatial frequencies and supra-threshold contrasts. However, the present work suggests that it is not sufficient to summarise ON and OFF pathway function using full-field stimuli, as we find the relationship between these two pathways interacts with polar angle.

The distribution of ON and OFF contrasts in the upper and lower visual field is currently unknown from the human perspective on natural scenes, and it is not trivial to investigate, considering that the upper and lower visual field are defined by gaze location, which varies with scene content and the passage of time. If the efficient transmission of information is the primary goal of the visual system, the difference in ON-OFF pathway tuning we have found between the UVF and LVF should be accompanied by a field-dependant ON-OFF contrast distribution. While this has not been previously considered in humans, it has been in the mouse. Using natural images taken from the Mouse perspective, Qiu et al. (2021) investigated the differences in dark bias between the UVF and LVF, finding that the dark bias was most prominent in the upper visual field. It is possible that this UVF bias reflects an adaptive specialisation to the mouse’s ecological niche, and it demonstrates that a polarity by visual field interaction can present in a terrestrial vertebrate. Future work should investigate the same question from the human perspective, where our data would predict the inverse to the mouse: an OFF dominance that is strongest in the LVF. Beyond the information theoretic approach, it is is possible that some contrast information is more behaviourally valuable, beyond an asymmetric scene distribution. That is, there is an ethological drive to the adaptive qualities of the human visual system, as has been suggested in mouse (Abballe & Asari, 2022).

### Implications for the assessment of ocular pathology

Most of the contrasts humans encounter in day-to-day life are supra-threshold (Balboa & Grzywacz, 2000; Cooper, 2016). Despite this, supra-threshold contrast perception is rarely investigated in pathological vision loss. For example, visual field perimetry focuses on incremental detection thresholds. This means we have a near absent quantitative understanding of the supra-threshold visual experience across the visual field of patients with ocular pathology. We have previously used the SSVEP to demonstrate an OFF-pathway vulnerability in Glaucoma (Norcia et al., 2022), but at a single contrast. Recently, Bham et al. (2020) measured behavioural contrast matching thresholds in patients with partial Glaucomatous field loss. At two contrast levels (2x and 4x patients’ absolute contrast threshold), they found that contrast matching was accurate, despite clear absolute threshold elevation. This means patients’ perception of contrast is preserved (and veridical) suprathreshold when the target and reference are identical in all spatial respects. From this limited evidence, it seems that Glaucoma patients do not experience an overall reduction in image contrast, which suggests the presence of a compensatory mechanism that allows contrast responses to “catch-up” beyond the absolute threshold for detection (but see Lek et al., 2019).

This “catch up” phenomenon is similar to loudness recruitment, a long-recognised consequence of hearing loss (Shi et al., 2022). It is possible that a similar phenomenon is at play in Glaucomatous visions loss, whereby a noise-limited detection mechanism is effected by retinal insult, while a suprathreshold mechanism remains relatively unaffected. This would manifest in a steepened contrast response function, and increasingly binarise percepts into clearly visible and entirely non-visible (as opposed to all stimuli becoming lower contrast as the disease progresses). The contrast response function is easily assessed using the SSVEP and relatable to discriminability, and direct neural measures of discriminability are possible (Nelson & Seiple, 1992). Furthermore, measuring the contrast response function using the SSVEP is a convenient way to objectively assess suprathreshold response slope at multiple visual field locations simultaneously. This could not only verify existing behavioural accounts using unsigned contrast, but can additionally bias responses towards particularly vulnerable processing pathways. The human ON- and OFF-pathways are thought to be differentially susceptible to Glaucomatous insult, with OFF retinal ganglion cells being particularly vulnerable (Norcia et al., 2022).

## Conclusion

Visual field asymmetries are present in the representation of incremental and decremental stimuli at multiple suprathreshold contrasts. These asymmetries may have their origins in the distribution of ON and OFF contrast in natural scenes and may be relevant for the assessment of suprthreshold vision in patients with visual-field dependent vision loss.

