## Supplemental Figures for "Visual field asymmetries in responses to ON and OFF pathway biasing stimuli"

### Supplementary material

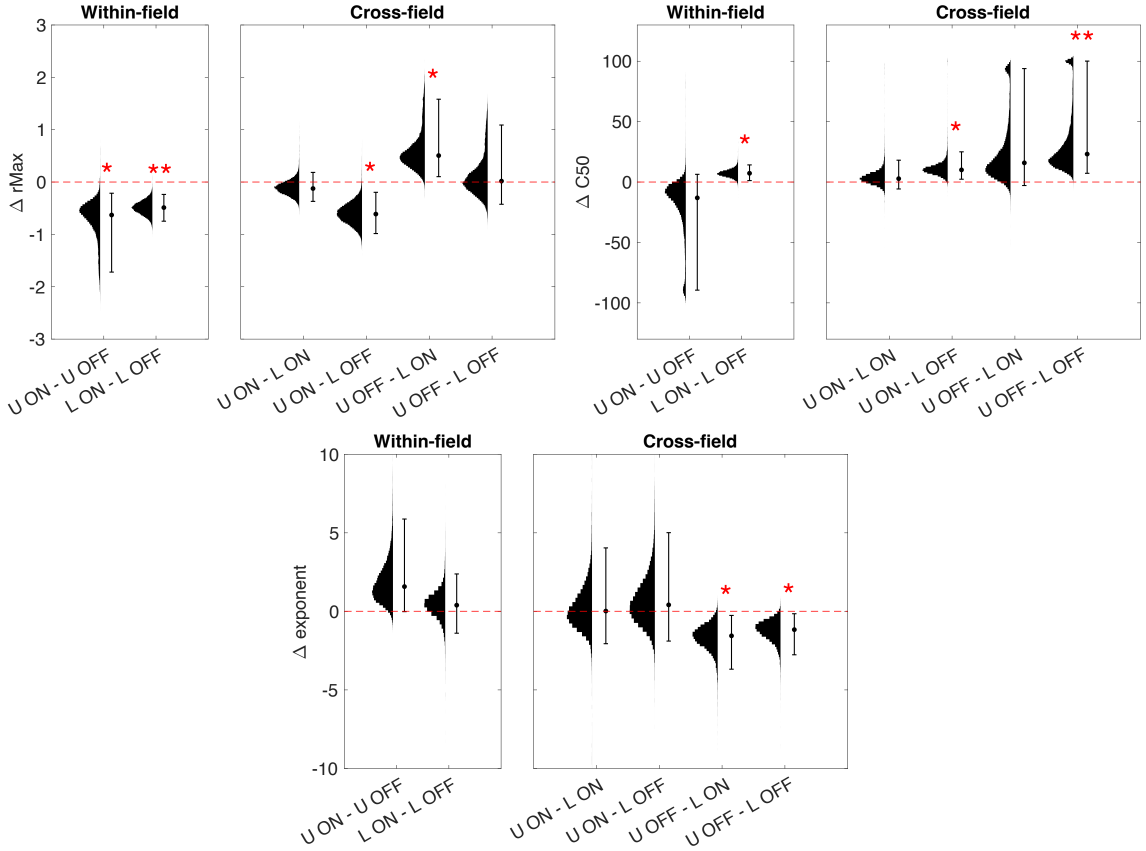


**Figure 9:** Half-violin plots showing the distributions yielded by condition-wise contrasts for RC1. The y-axis label denotes the parameter being contrasted, while the x-axis labels denote the conditions entered into the contrast. Each parameter is split by within-field comparisons and cross-field comparisons. Significant differences are indicated by red asterisks (* = <.05, ** < .01)

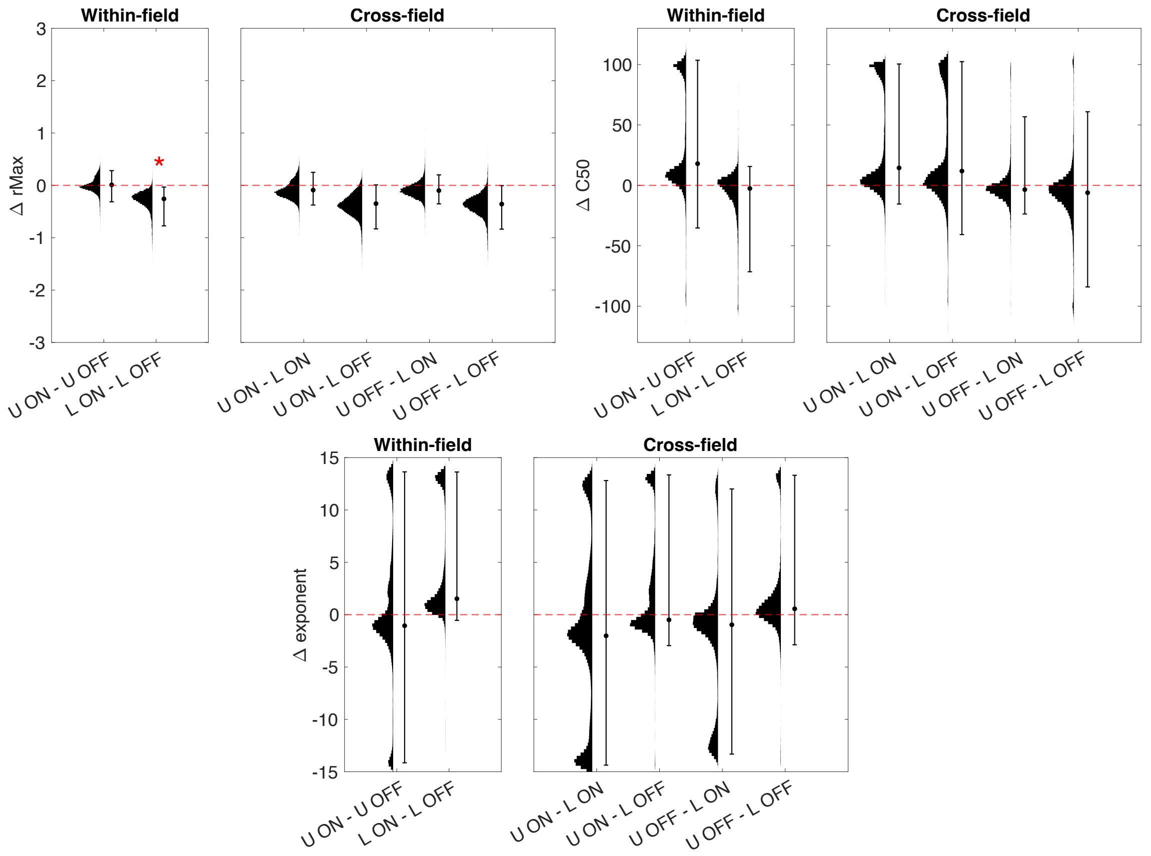


**Figure 10:** Half-violin plots showing the distributions yielded by condition-wise contrasts for RC2. The y-axis label denotes the parameter being contrasted, while the x-axis labels denote the conditions entered into the contrast. Each parameter is split by within-field comparisons and cross-field comparisons. Significant differences are indicated by red asterisks (* = <.05, ** < .01)
